# Identification of the safe harbor locus, AAVS1, from porcine genome and site-specific integration of recombinase-mediated cassette exchange system in porcine fibroblasts using CRISPR/Cas9

**DOI:** 10.1101/2021.09.15.460532

**Authors:** Choongil Lee, Soo-Young Yum, Woojae Choi, Seokjoong Kim, Goo Jang, Okjae Koo

## Abstract

Gene integration at site-specific loci, such as safe harbor regions for stable expression via transgenesis, is a critical approach for understanding the function of a gene in cells or animals. The AAVS1 locus is a well-known safe harbor site for human and mouse studies. In the present study, we found an AAVS1-like sequence in the porcine genome using the UCSC Genome Browser and designed TALEN and CRISPR/Cas9 to target AAVS1. The efficiency of CRISPR/Cas9 for targeting the AAVS1 locus in porcine cells was superior to that of TALEN. An AAVS1-targeting donor vector containing GFP was designed and cloned. We added a loxP-lox2272 cassette sequence to the donor vector for further exchange of various transgenes in the AAVS1-targeted cell line. The donor vector and CRISPR/Cas9 components targeting AAVS1 were transfected into a porcine fibroblast cell line. Targeted cells of CRISPR/Cas9-mediated homologous recombination were identified by antibiotic selection. Gene knock-in at the AAVS1 locus was confirmed by PCR analysis. To induce recombinase-mediated cassette exchange (RMCE), another donor vector containing the loxP-lox2272 cassette and inducible (Tet-on) Cre recombinase was cloned. The Cre-donor vector was transfected into the AAVS1-targeted cell line, and RMCE was induced by adding doxycycline to the culture medium. RMCE in porcine fibroblasts was confirmed using PCR analysis. In conclusion, gene targeting at the AAVS1 locus and RMCE in porcine fibroblasts was successful. This technology will be useful for future porcine transgenesis studies and the generation of stable transgenic pigs.

## 1. Introduction

Pigs are valuable models for human diseases because they have many similarities to humans, such as organ size and physiology [1]. On the other hand, pigs are regarded as the most suitable xenotransplantation donor animal for human diseases [2]. For these purposes, various genetic engineering techniques, such as overexpression [3], conditional expression [4], knock-out (KO) [5,6], and knock-in (KI) have been developed in pigs. Recently, the development of genome engineering tools such as zinc-finger nuclease (ZFN), transcription activator-like (TAL) effector nucleases (TALEN), and clustered regularly interspaced short palindromic repeat/Cas9 (CRISPR/Cas9), have made it possible to modify the porcine genome more efficiently [5–7].

One of the most important objectives of transgene expression is targeted disruption and insertion of a gene of interest into the target site for stable gene expression without harming the organism. For this purpose, genomic safe harbors and unique genome sites that do not affect the survival of an organism and do not influence the expression of surrounding genomic regions have been investigated [8]. Thus far, well-known safe harbor regions are *ROSA26* [9], *CCR5* [10], and *AAVS1* [11] in human and mouse genomes. Furthermore, various genomic studies have been conducted on these regions.

*AAVS1* is not only an adenovirus integration site in various animal genomes, including that of humans [12], but also a safe harbor region for gene insertion [13–15]. Recently, *ROSA26* knock-in and *H11* knock-in pig models have been generated by safe harbor integration using CRISPR/Cas9 [16]. The porcine *AAVS1* locus (p*AAVS1)* is another valuable safe harbor for generating pig models by gene editing; however, the p*AAVS1* locus has not yet been identified. In the current study, we identified the p*AAVS1* locus in the porcine genome. In addition, we knock-in a recombinase-mediated cassette exchange system into p*AAVS1* in porcine fibroblasts using CRISPR/Cas9 technology. This cell line can be used for further transgenic studies in porcine models.

## 2. Materials and Methods

### 2.1. Identification of pAAVS1 using bioinformatics tools

Human *PPP1R12C* gene Exon1/2 sequences (S1 Appendix) were used to search for homologous sequences in the pig genome. Using BLAT in the UCSC Genome Browser (https://genome.ucsc.edu/index.html), we found an overlapping DNA sequence, *PPP1R12C* (ENSSSCG00000027189 & NC_010448.3 Chromosome 6 Reference Sscrofa10.2 Primary Assembly), on chromosome 6 in the pig genome.

### 2.2. RT-PCR of porcine PPP1R12C

Total RNA was extracted to analyze the gene expression in each of the porcine organs (liver, lung, spleen, heart, kidney, testis, and ovary) and cultured porcine fibroblast cells using Trizol reagent (Invitrogen). Complementary DNAs (cDNAs) were synthesized using Maxime RT Premix (iNtRON) according to the manufacturer’s protocol. Information on the primers used is listed in Supplementary Table 1. The gene expression of *PPP1R12C* and *GAPDH* was measured using ImageJ (NIH).

### 2.3. Cell culture

All TALEN and CRISPR/Cas9 plasmids were obtained from a commercial company (ToolGen). We transfected 1 × 10^6^ immortalized cells using the Neon nucleofection system and 10 μg of plasmid DNA at a weight ratio of 25:25:50 (plasmid encoding Cas9: plasmid encoding sgRNA: plasmid for HDR vector) according to the manufacturer’s protocol. The transfected cells were cultured for 2 days at 37 °C and subjected to blasticidin S hydrochloride for 1 week.

### 2.4. T7E1 assay and KO sequencing

Genomic DNA was extracted using the G-DEX IIc Genomic DNA Extraction Kit (iNtRON) after 3 days of transfection. TALEN target sites were PCR-amplified using the primer pairs listed in Supplementary Table 1. To analyze the genome editing efficiency of TALEN and CRISPR/Cas9 used in this study, T7E1 analysis was performed. Briefly, the amplicons were denatured by heating and annealed to form heteroduplex DNA, which was treated with 5 units of T7 endonuclease 1 (New England Biolabs) for 20 min at 37 °C and analyzed by 2% agarose gel electrophoresis. The sequence of the amplicons was analyzed by a commercial company (Marcrogen).

### 2.5. KI and PCR sequencing

The Tet-on vector was obtained from the Lenti-X Tet-One Inducible Expression System (Puro) (Clontech, Cat. No. 634847). We transfected 1 × 10^6^ HDR-immortalized cells using the Neon nucleofection system and 10 μg of plasmid DNA (plasmid encoding Cre recombinase, cassette exchange vector flanked by loxP and lox2272) according to the manufacturer’s protocol. The transfected cells were cultured for 1 week at 37 °C. During culture, cells were treated with doxycycline on the 1st day and puromycin from the 2nd day onwards.

## 3. Results and Discussion

### 3.1. Identification of pAAVS1 DNA sequence in pig genome

To identify the *AAVS1* DNA sequence in the pig genome, we initially performed a homology search using BLAT in the UCSC Genome Browser. By comparing the human and pig genomes, protein phosphatase 1 regulatory subunit 12C (*PPP1R12C*) (ENSSSCG00000027189 & NC_010448.3 Chromosome 6 Reference Sscrofa10.2 Primary Assembly) was identified as a potential *AAVS1* DNA sequence in the pig genome. To consider the *PPP1R12C* locus as a safe harbor for ectopic gene insertion, we determined the expression level of *PPP1R12C* mRNA in various pig organs. *PPP1R12C* gene expression in specific organs (liver, lung, spleen, heart, kidney, ovary, and testis) was observed, and it was stably expressed in all the tested organs (Fig 1). Therefore, the porcine *PPP1R12C* locus is considered a promising candidate for porcine gene editing. To modify the gene, immortalized porcine fibroblast (iPF) cells, which have been used for gene editing in a previous study [17], were used in this study.

**Fig 1.**
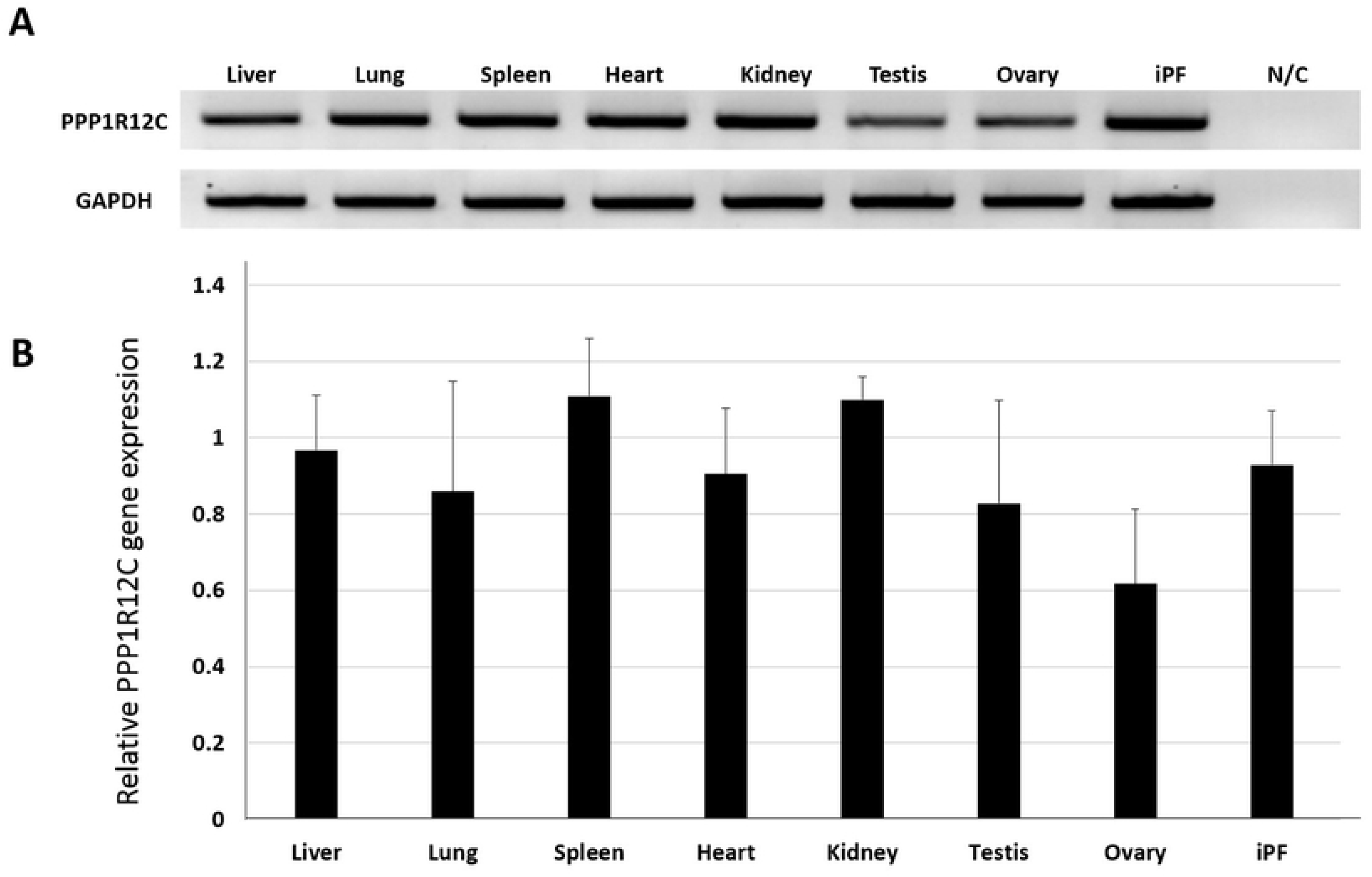
*PPP1R12C* expression in various porcine organs. A. Organ-specific *PPP1R12C* expression. B. Relative *PPP1R12C* expression level.

### 3.2. PPP1R12C gene modification by TALEN and CRISPR/Cas9

For efficient gene modification, two types of endonucleases, TALEN and CRISPR/Cas9, were designed. Among the designed endonucleases, the target cleavage activity of CRISPR/Cas9 was better than that of TALEN for *PPP1R12C* gene modification (Fig 2A). CRISPR/Cas9-mediated mutations were confirmed by DNA sequencing analysis (Fig 2B). The DNA mutation induced frameshifts (1 bp insertion, 1 bp deletion, and 16 bp deletion) and large deletions (30 bp deletion; 10 amino acids removed) in the sequence.

**Fig 2.**
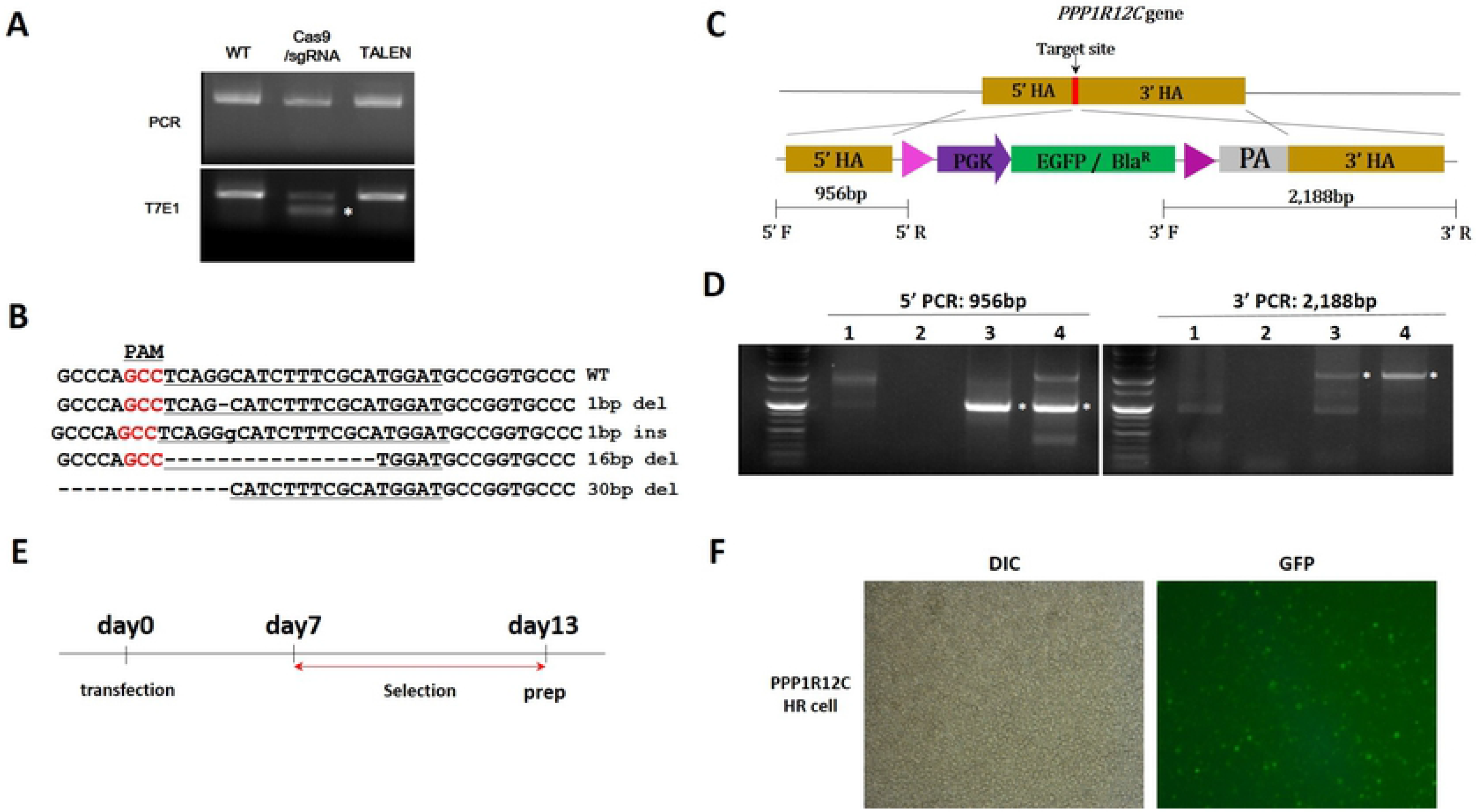
Gene modification of the *PPP1R12C* locus. A. T7E1 assay of Cas9/sgRNA and TALEN activity at the *PPP1R12C* site. B. Mutated DNA sequences of the AAVS1-like genome region after Cas9/sgRNA treatment. C. HDR scheme of the *PPP1R12C* site. D. HDR results were confirmed by PCR. 1: Cas9/sgRNA, 2: donor vector, 3: before drug selection, 4: after drug selection. E. Experimental step for selection of HDR-positive cells. Cells were selected by 20 μg/ml of blasticidin S hydrochloride for 1 week. F. Fluorescence image of HDR-positive cells. White asterisk denotes the correct DNA band size.

### 3.3. Homology-directed repair using PPP1R12C gene as a safe-harbor locus

After confirmation of CRISPR/Cas9 activity, we designed a homology-directed repair (HDR) vector for KI using the *PPP1R12C* site as a safe harbor (Fig 2C). This HDR vector contained 683 bp of a 3 homology arm with AAVS1_sgRNA target sequences and 2050 bp of a 5′ homology arm [18], EGFP and blasticidin S hydrochloride resistance coding sequences as a selectable marker, and loxP/lox2272 sequences for Cre recombinase/loxP-mediated cassette exchange in further experiments. iPF cells were transfected with Cas9, *PPP1R12C* sgRNA, and an HDR donor vector. After 13 days of transfection, HDR results were confirmed by PCR analysis before and after blasticidin S hydrochloride selection (Fig. 2D, E).

To detect homologous recombination-mediated modifications, four pairs of primers (upstream and downstream) were designed. For the upstream primer pair, one was localized outside the left homologous arm to exclude random insertion, and the other was localized inside the loxP sequence. For the downstream primer pair, one was localized outside the right homologous arm, and the other was localized downstream of the gene encoding blasticidin S hydrochloride. All PCR products were sub-cloned and sequenced to characterize the modifications (Fig S1, S2). Moreover, *PPP1R12C*-mediated HDR gene insertion was confirmed by the stable expression of *EGFP* (Fig 2F). Therefore, *PPP1R12C* can serve as a safe harbor in the pig genome.

### 3.4. Recombinase-mediated cassette exchange using PPP1R12C mediated HDR cells

To induce recombinase-mediated cassette exchange (RMCE), we designed a Cre recombinase/loxP-mediated cassette exchange vector for use in the gene of interest in the pig genome (Fig 3A). To detect cassette exchange-mediated modification, four pairs of primers (upstream, downstream, and two internal regions) were designed. For the upstream primer pair, one was localized to the left homologous sequence and the other was localized inside the SV40 poly-A tail gene. For the downstream primer pair, one was localized on the right homologous arm, and the other was localized on the WPRE. Cre/loxP-mediated cassette exchange results were confirmed by PCR analysis after blasticidin S hydrochloride selection (Fig 3B, C), and DNA sequences were analyzed by sequencing (Fig S3, S4). GFP-negative cells (Fig 3D) were detected, and the results were consistent with the PCR results. Therefore, gene modification can be performed using *PP1R12C* as a safe harbor.

**Fig 3.**
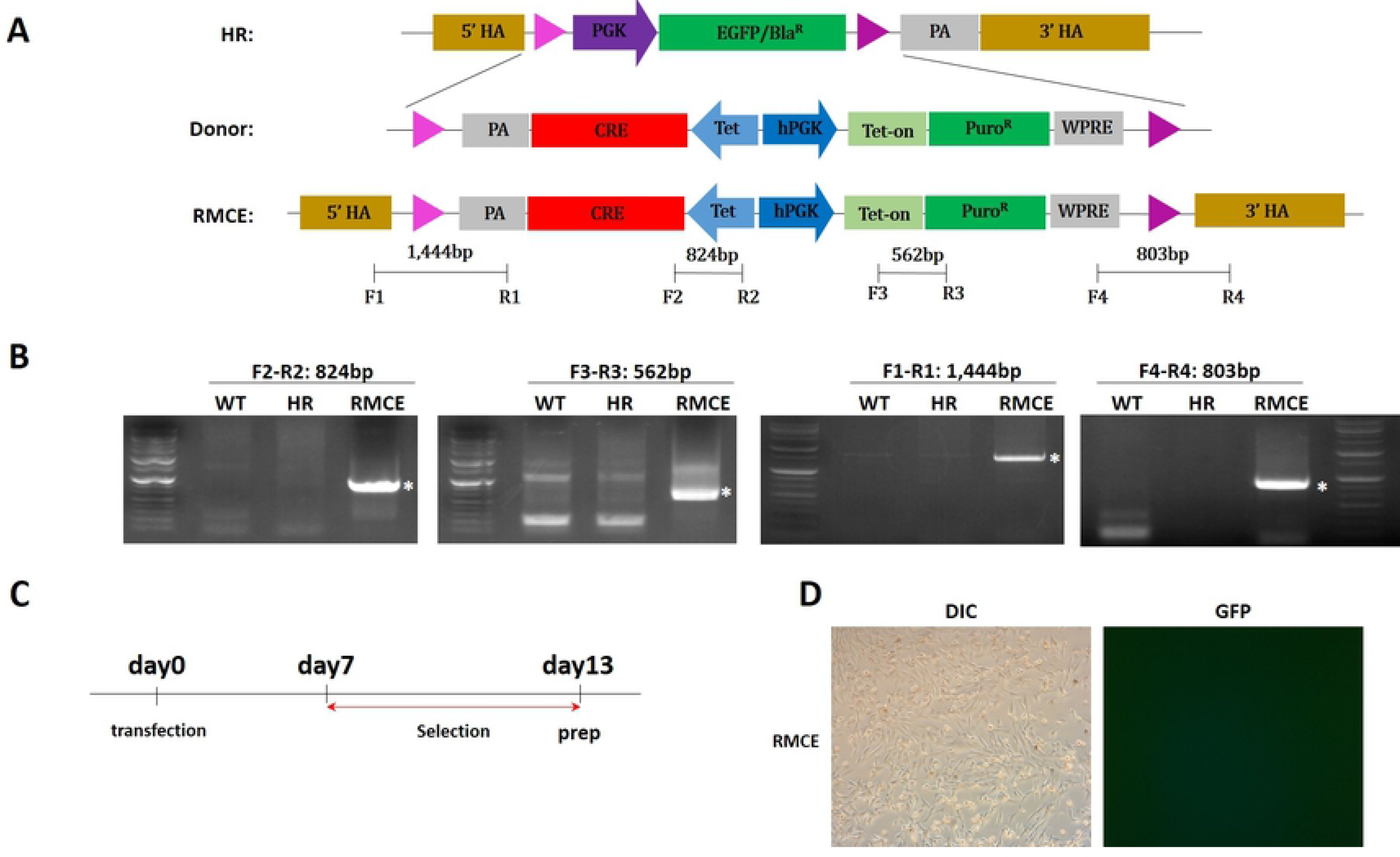
Cre/loxP-mediated cassette exchange A. Cre/loxP-mediated cassette exchange scheme. B. Cassette exchange results were confirmed by PCR. C. Experimental step for selection of Cre/loxP-mediated cassette exchange-positive cells. Cells were selected by 10 μg/ml of puromycin for 1 week. D. Fluorescence image of cassette-exchanged cells. White asterisk denotes the correct DNA band size.

## 4. Conclusions

In conclusion, the data demonstrated that the *PPP1R12C* locus is a good candidate as a safe harbor for knock-in experiments in pig models. In future studies, knock-in experiments using *PPP1R12C* of the pig genome would result in stable gene expression.

## Acknowledgments

This study was financially supported by the Rural Development Administration (Project No. PJ009802), Korea.

---

**S1 Appendix.** Human PPP1R12C gene Exon1/2 sequences.

**Fig S1.** HR is confirmed by PCR (5’: outside of 5’ homology arm - loxP).

**Fig S2.** HR is confirmed by PCR (3’: EGFP-Blasticidin R - outside of 3’ homology arm).

**Fig S3.** Cassette exchange is confirmed by PCR (up-stream: 5’ homology arm & CRE recombinase).

**Fig S4.** Cassette exchange is confirmed by PCR (down-stream: WPRE & 3’homology arm).

**Table S1.** Sequences of each primer.

## Notes

### Competing Interest Statement

The authors have declared no competing interest.

## References

1. Sachs DH. The pig as a potential xenograft donor. Vet Immunol Immunopathol. 1994;43: 185–191. doi:10.1016/0165-2427(94)90135-x

2. Yao J, Huang JJ, Zhao JG. Genome editing revolutionize the creation of genetically modified pigs for modeling human diseases. Hum Genet. 2016;135: 1093–1105. doi:10.1007/s00439-016-1710-6

3. Lai LX, Kang JX, Li RF, Wang JD, Witt WT, Yong HY, et al. Generation of cloned transgenic pigs rich in omega-3 fatty acids. Nat Biotechnol. 2006;24: 435–436. doi:10.1038/nbt1198

4. Li S, Flisikowska T, Kurome M, Zakhartchenko V, Kessler B, Saur D, et al. Dual Fluorescent Reporter Pig for Cre Recombination: Transgene Placement at the ROSA26 Locus. PLoS One. 2014;9: 8. doi:10.1371/journal.pone.0102455

5. Carlson DF, Tan WF, Lillico SG, Stverakova D, Proudfoot C, Christian M, et al. Efficient TALEN-mediated gene knockout in livestock. Proc Natl Acad Sci U S A. 2012;109: 17382–17387. doi:10.1073/pnas.1211446109

6. Hauschild J, Petersen B, Santiago Y, Queisser AL, Carnwath JW, Lucas-Hahn A, et al. Efficient generation of a biallelic knockout in pigs using zinc-finger nucleases. Proc Natl Acad Sci U S A. 2011;108: 12013–12017. doi:10.1073/pnas.1106422108

7. Yang LH, Guell M, Niu D, George H, Lesha E, Grishin D, et al. Genome-wide inactivation of porcine endogenous retroviruses (PERVs). Science. 2015;350: 1101–1104. doi:10.1126/science.aad1191

8. DeKelver RC, Choi VM, Moehle EA, Paschon DE, Hockemeyer D, Meijsing SH, et al. Functional genomics, proteomics, and regulatory DNA analysis in isogenic settings using zinc finger nuclease-driven transgenesis into a safe harbor locus in the human genome. Genome Res. 2010;20: 1133–1142. doi:10.1101/gr.106773.110

9. Soriano P. Generalized lacZ expression with the ROSA26 Cre reporter strain. Nat Genet. 1999;21: 70–71. doi:10.1038/5007

10. Perez EE, Wang JB, Miller JC, Jouvenot Y, Kim KA, Liu O, et al. Establishment of HIV-1 resistance in CD4(+) T cells by genome editing using zinc-finger nucleases. Nat Biotechnol. 2008;26: 808–816. doi:10.1038/nbt1410

11. Smith JR, Maguire S, Davis LA, Alexander M, Yang FT, Chandran S, et al. Robust, persistent transgene expression in human embryonic stem cells is achieved with AAVS1-targeted integration. Stem Cells. 2008;26: 496–504. doi:10.1634/stemcells.2007-0039

12. Linden RM, Ward P, Giraud C, Winocour E, Berns KI. Site-specific integration by adeno-associated virus. Proc Natl Acad Sci U S A. 1996;93: 11288–11294. doi:10.1073/pnas.93.21.11288

13. Orlando SJ, Santiago Y, DeKelver RC, Freyvert Y, Boydston EA, Moehle EA, et al. Zinc-finger nuclease-driven targeted integration into mammalian genomes using donors with limited chromosomal homology. Nucleic Acids Res. 2010;38: 15. doi:10.1093/nar/gkq512

14. Hockemeyer D, Soldner F, Beard C, Gao Q, Mitalipova M, DeKelver RC, et al. Efficient targeting of expressed and silent genes in human ESCs and iPSCs using zinc-finger nucleases. Nat Biotechnol. 2009;27: 851–U110. doi:10.1038/nbt.1562

15. Maresca M, Lin VG, Guo N, Yang Y. Obligate Ligation-Gated Recombination (ObLiGaRe): Custom-designed nuclease-mediated targeted integration through nonhomologous end joining. Genome Res. 2013;23: 539–546. doi:10.1101/gr.145441.112

16. Ruan JX, Li HG, Xu K, Wu TW, Wei JL, Zhou R, et al. Highly efficient CRISPR/Cas9-mediated transgene knockin at the H11 locus in pigs. Sci Rep. 2015;5: 10. doi:10.1038/srep14253

17. Moon J, Lee C, Kim SJ, Choi JY, Lee BC, Kim JS, et al. Production of CMAH Knockout Preimplantation Embryos Derived From Immortalized Porcine Cells Via TALE Nucleases. Mol. Ther. Nucleic Acids. 2014;3: 8. doi:10.1038/mtna.2014.15

18. Shin J, Chen JK, Solnica-Krezel L. Efficient homologous recombination-mediated genome engineering in zebrafish using TALE nucleases. Development. 2014;141: 3807–3818. doi:10.1242/dev.108019

